# Neonatal mimicry of caregivers at home: feasibility of an asynchronous online paradigm

**DOI:** 10.1101/2025.06.26.661799

**Authors:** Katherine Casey, Benazir Neree, Camila Larrauri, Guangyu Zeng, Kira Ashton, Jeff Gill, Elizabeth A. Simpson, Laurie Bayet

## Abstract

While early life sets the stage for later learning, comparatively less is known about newborns’ cognition than that of older infants. A striking example is the lack of consensus regarding the extent to which newborns spontaneously mimic gestures, and whether such behavior drives bonding and learning. Despite the theoretical importance of these questions, practical challenges limit researchers’ ability to engage newborns in behavioral research. Webcam-based, asynchronous online studies have expanded developmental science’s capacity to reach older infants. However, such scalable and replicable methods have yet to be deployed with younger infants. Taking a commonly-used neonatal mimicry paradigm as a test case, we assessed the feasibility of leveraging an open-source online platform (Children Helping Science) for asynchronous research with 0-6-week-olds and their caregivers. Caregivers modeled face movements to their 4-45 days-old infants (N = 29, N = 17 included) while webcams filmed their infants’ responses; 13 dyads participated more than once (72 included test videos). There was preliminary, moderate evidence against group-level neonatal mimicry of caregivers’ tongue protrusions (Bayes Factor ∼ ⅓), and inconclusive evidence for or against mimicry of caregivers’ mouth openings (⅓ < BF < 3). Simulations identified a target sample size for more conclusive evidence. Finally, we asked whether caregivers perceived their newborns’ behavior as imitative. Caregivers’ perceptions of mimicry reflected infants’ behaviors but did not align with an often-used metric of mimicry (“imitators”). These results demonstrate the feasibility of asynchronous online behavioral studies with newborns and provide a foundation for future research on neonatal mimicry of caregivers.x1

---

The first weeks of life build a critical foundation for later health and learning. As their brains and minds undergo rapid development, young infants already engage with the world as budding cognitive and social individuals; therefore, a better understanding of their abilities could facilitate the delivery of very early screening or interventions (Guellai & Streri, 2022; Streri et al., 2013). Despite the importance of this developmental period, infants within the first 2 months of life remain relatively under-studied by current behavioral research, limiting the field’s ability to settle theoretical debates or inform interventions (Nagy, 2011). This gap is problematic for developmental science because it prevents the field from precisely delineating *what* develops in early postnatal life, i.e., what the starting point of postnatal development is, at birth. For example, the existence of neonatal mimicry —whether or not newborns and young infants spontaneously mimic gestures in face-to-face interactions— continues to generate a remarkably great deal of controversy, despite its theoretical significance and having been first reported nearly 50 years ago (Meltzoff & Moore, 1977). To advance a new approach that may help to address these long-standing issues, the current study leveraged an existing open-source online platform (Children Helping Science, CHS, formerly known as Lookit; Scott et al., 2017; Scott & Schulz, 2017) to investigate the feasibility of asynchronously engaging young infants and their caregivers in behavioral research from their homes, and thereby inform the design of larger studies.

Practical challenges inherent to infant behavioral studies, such as recruitment challenges, shorter attention spans, higher attrition, and consequently smaller sample sizes (Coyne, 2016), are only magnified in the neonatal period. Scheduled daytime testing (including scheduled remote testing) inherently struggles to accommodate young infants’ variable arousal states and immature circadian rhythms (Rivkees, 2003). The resulting paucity of usable newborn behavioral data contributes to enduring debates in our understanding of what young infants’ minds can or cannot do (for a recent example, Aslin et al., 2023; Blumberg & Adolph, 2023; Liu et al., 2023). The CHS platform allows families to participate in behavioral studies online at any time, using only their own computer and webcam, without a researcher present. Such asynchronous online research delivers studies that are intrinsically more easily scalable and replicable, are less burdensome to participants, and can recruit more representative samples (Scott et al., 2017; Scott & Schulz, 2017). Yet, while asynchronous online data collection is feasible and reliable with infants as young as 4-5 months in looking time paradigms (Chuey et al., 2024; Scott et al., 2017; Sheskin et al., 2020), there is little data to date about testing younger infants online—including in neonatal mimicry paradigms.

Neonatal mimicry, sometimes called neonatal imitation, refers to the potential capacity of newborns and young infants to reproduce or match someone else’s gestures, including facial and hand movements (Davis et al., 2021; Jones, 2017; Oostenbroek et al., 2013; Simpson et al., 2014). Multiple studies have reported mimicry in 0-6 week-old infants, most often of tongue protrusion (TP), mouth opening (MO), or finger movement (Meltzoff & Moore, 1977, 1983; Nagy et al., 2005, 2013, 2014, 2020). However, multiple studies have also reported null or mixed effects at the group level (Anisfeld, 1991; Anisfeld et al., 2001; Jacobson, 1979; Meltzoff et al., 2018; Oostenbroek et al., 2016). Concerningly, meta-analytic evidence suggests that researcher affiliation significantly moderates the size of published neonatal mimicry effects, over and beyond certain methodological choices such as test duration (Davis et al., 2021). This striking finding suggests that either some other methodological choices or experimenter bias (Davis et al., 2021) (in either direction) may underlie some of the inconsistencies in the neonatal mimicry literature (see also Meltzoff et al., 2019). Therefore, transparent paradigms that can be deployed by multiple laboratories may be necessary to resolve the debates surrounding neonatal mimicry, over and beyond meta-analyses.

The question of the existence and prevalence of neonatal mimicry carries far-reaching implications for theories of social learning and face processing. Early imitation plays a foundational role in some developmental theories (Meltzoff & Decety, 2003; Trevarthen & Aitken, 2001), yet its genuine occurrence would directly falsify others (Heyes, 2016; Jones, 2017; Keven & Akins, 2017; Ray & Heyes, 2011). While the current study does not definitively adjudicate between these theories, some further implications are briefly outlined below to illustrate the conceptual importance of neonatal mimicry. The true presence of neonatal mimicry, demonstrated with sufficient control conditions to rule out alternative interpretations such as arousal (Anisfeld, 1991; Keven & Akins, 2017), and occurring at a sufficient prevalence to be detectable at the group level, would imply that imitation does not require extensive social learning or maturation (Guellai & Streri, 2022; McGowan & Delafield-Butt, 2023). Neonatal mimicry of *facial* movements would additionally imply that newborns cross-modally map the visual representation of their caregiver’s face with the proprioceptive-motor output of their own facial motor movements, though both representations remain immature (Anisfeld, 1996; Meltzoff, 2007; Simpson et al., 2014). If so, then proprioceptive-motor feedback could theoretically scaffold visual learning of other people’s face movements from birth, when visual acuity remains poor. Indeed, data from adults with Moebius syndrome, a condition that affects cranial nerves and causes paralysis of facial muscles, suggest that motor feedback significantly contributes to recognizing facial movements in adulthood (Japee et al., 2023). For such potentially far-reaching developmental impacts to be realized, however, neonatal mimicry needs to occur in interactions with caregivers. Conversely, the true absence of neonatal mimicry at the group level would provide support for either of two alternatives. One possibility is that young infants’ visual and/or motor systems may not yet be mature enough to represent others’ dynamic face movements or map them cross-modally from birth. If so, then young infants’ processing of other people’s face movements may initially be restricted to the visual modality. An alternative possibility is that infants represent face movements cross-modally from birth, but most do not spontaneously or robustly mimic these movements until later in development. Nonetheless, some individual newborns may spontaneously mimic others (Heimann & Schaller, 1985; Kaburu et al., 2016; Simpson et al., 2016), or genuine instances of neonatal mimicry could happen sporadically, with potential impacts on individual development.

Thus, beyond the question of whether neonatal mimicry can be evoked under carefully-controlled laboratory conditions, a separate question is whether newborns and young infants generally tend to mimic their caregivers’ gestures under real-world conditions. Infants might be more inclined to mimic in favorable contexts or behavioral states, such as at home with their caregivers. More generally, caregiver-infant behavior in the naturalistic home environment may slightly differ from what can be observed in the laboratory (Belsky, 1980; Lamm et al., 2014). In addition, most studies use researchers rather than caregivers as models, to increase experimental control and safeguard against the possibility that any imitative responses could be learned (Meltzoff & Moore, 1983).

For example, of the 26 neonatal mimicry studies reviewed in Davis et al. (2021), only 3 reported using the mother as the model (Heimann & Schaller, 1985; Meltzoff & Moore, 1992; Ullstadius, 1998), and of those, only 1 was conducted in infants’ most ecological environment, i.e., at home (Heimann & Schaller, 1985). None reported using non-maternal caregivers, such as fathers, as the model, although older infants engage in imitation games with fathers as well as with mothers (Kokkinaki & Kugiumutzakis, 2000). In the United States, most of young infants’ face-to-face interactions occur with only a few (e.g., 1-3) people, such as primary caregivers (Jayaraman et al., 2015; Sugden et al., 2014). Therefore, to directly support early social learning or bonding (Bjorklund, 1987; Simpson et al., 2014; Užgiris, 1991), neonatal mimicry must occur within those crucial interactions with caregivers, their earliest social partners.

Even in the absence of any genuine neonatal mimicry (e.g., if the phenomenon is artefactual ; Anisfeld, 1991; Keven & Akins, 2017), caregivers’ *perception* that their child is attempting to mimic them may still indirectly boost early learning and bonding. For example, thinking that their young infant child is trying to mimic them may boost caregivers’ mind-mindedness (McMahon & Bernier, 2017), attunement (Kokkinaki, 2003), positive affect, or motivation to engage in face-to-face interactions or imitative games (Markodimitraki & Kalpidou, 2021). Yet, few studies have examined whether and when caregivers perceive their infants’ responses as mimicry (Heimann & Schaller, 1985). More generally, relatively little is known about which infant behaviors are *perceived* as communicative or indicative of a desire for social connection by caregivers themselves, as opposed to developmental researchers (Akhtar & Jaswal, 2020).

To address these gaps, the current study used the CHS asynchronous online platform to investigate young infants’ mimicry of caregivers’ face movements and caregivers’ perceptions of mimicry. Caregivers were randomly assigned to model either tongue protrusion (TP) or mouth opening (MO) to their infant, two facial gestures most often used in neonatal mimicry studies (Davis et al., 2021). Dyads could participate multiple times, yielding partially longitudinal data. The study first aimed to establish the feasibility of engaging 0-6-week-olds and their caregivers in asynchronous behavioral studies online, which would ultimately allow for larger sample sizes. We assessed paradigm completion, attrition, and demographic representativity. To promote transparency and facilitate replication, paradigm materials are shared. The study additionally aimed to obtain preliminary estimates of the extent to which young infants mimic facial gestures from their caregivers, and the extent to which caregivers interpret their child’s responses during the task as imitative.

## 1. Materials and Methods

### 1.1. Participants

A total of 29 full-term infants (4-45 days-old, born at or later than 38 weeks of gestational age, 16 female) and caregivers from the United States participated on the CHS platform. Infants’ ages covered the typical range of neonatal mimicry studies, i.e., approximately 6 weeks old or younger (Davis et al., 2021). In total, 129 study sessions from these N = 29 dyads (N = 17 included) were run, resulting in 118 test trial videos (72 included). Participants could complete a study session more than once and were compensated $15 for each, up to once daily until the maximal inclusion age of 45 days. Almost half of participants (n = 13, 45%) completed more than one session (average: n = 4.45 ± 7.34 sessions by participant, range 1-34). Infants were an average of 23.24 days old on their first session (N = 29, range: 4-41 days), and 23.71 days old on their first session for included infants only (N = 17, range 4-41 days). Participants were recruited through online advertisements, CHS, and Prolific. An additional 6 sessions were recorded but not considered further due to the video not showing a child, or the participating child being older than 45 days.

Participating caregivers recorded themselves providing verbal informed consent at the start of each online session. Procedures were conducted according to ethical standards and approved by the Institutional Review Board (IRB) of [masked]. Data were collected in approximately 18 weeks, spanning from December 2020 to April 2021. The total sample size was determined by funding constraints. To be eligible for compensation, participating study sessions had to include a valid consent statement, and participating infants had to be within the included age range and be visible on the video with their caregiver during the test trial.

### 1.2. Paradigm and stimuli

Participating caregivers provided demographic information, including their infant’s birth date, their own age, and other family characteristics (**Supplementary Methods**). Caregivers were randomly assigned to model either TP or MO and were guided through a 180 second burst-pause paradigm (Meltzoff & Moore, 1983) by pre-recorded auditory cues. The paradigm consisted of a 20 second still-face period, followed by a 20 second gesture period (either TP or MO), repeated to mimic natural infant-caregiver turn-taking interactions, for a total duration of 180 seconds (**Supplementary Table 1**). To avoid confusing caregivers, minimize attrition, and avoid within-subject spillover across conditions, we used a between-subjects design. Caregivers were randomly assigned to either condition (TP or MO) on their first session and stayed in that same condition in subsequent sessions (if any). Due to a technical error, one participant contributed one included session in a different condition (MO) than their assigned condition (TP). This session (the participant’s first included session) was excluded from participant-level, between-participants analyses (i.e., **Results** section 2.3.1) but remained included in all other (session-level) analyses.

Prior to launching the test trial, participating caregivers saw a video clip demonstrating the facial movement they had been assigned to model to their child (see **Data Availability**), and pictures demonstrating proper positioning with regards to the webcam. Then, they recorded themselves practicing the target movements on their own in front of the camera (**Supplementary Figure 1A**). After practice, caregivers started the test trial, in which they completed the mimicry paradigm with their infant. During the 180s test trial, participants heard auditory cues guiding them to either model a resting face (“Rest, continue looking at your baby, but make a neutral face”) or model the target gesture (“Go, continue looking at your baby, and [stick out your tongue/open your mouth]”) in 20s segments (**Supplementary Table 1** and **Supplementary Figure 1B**). These auditory cues were matched in length (4s) and volume (peak dB) between the TP and MO conditions in Audacity, and enabled the caregivers to maintain eye contact with their infants for the duration of the test without having to look up at their screens.

To better accommodate young infants’ labile arousal states, caregivers could stop and restart the test trial within the same session as needed based on their infant’s state (e.g., hungry, sleepy, fussy). In such cases (23 out of 129 sessions), the most complete test period and the earliest usable baseline period within that same test session (e.g., first resting-face period before gesture modeling began) were coded for infant gestures and used in analyses.

After completing the test trial, participating caregivers reported whether they thought that their infant was trying to make the gesture back at them at least some of the time during the trial. Caregivers first provided a “forced choice” binary response, i.e., to “If you were to guess, would you say that your child was trying to [stick out their tongue/open their mouth] back at you at least some of the time when you were making that gesture? Just guess if you are not sure”, they chose “Yes, my child was trying to make the face back at me” or “No, my child was not trying to make the face back at me”. Then, they reported the degree to which they felt confident in their answer on a scale of 0-5.

To further characterize the representativeness of the sample with regards to post-partum depressive symptoms, caregivers completed the Participant Health Questionnaire (PHQ-9) survey (Kroenke et al., 2001). Caregivers were provided with links to online resources if their PHQ-9 score met or exceeded a clinical threshold of 10 (Flynn et al., 2011). If they completed the PHQ-9 several times across repeated sessions, we retained only their maximum PHQ-9 score.

### 1.3. Video preprocessing

Each test trial video was pre-processed twice using Adobe Premiere Pro, in two parallel streams to be used by two coder teams. In the *first* stream, the *infant*’s face was (Gaussian) blurred, leaving the caregiver’s face and condition-specific audio cues intact (**Supplementary Figure 1C**). Along with the “practice” videos (**Supplementary Figure 1A**), these videos were used for the first set of coders to check video quality and task fidelity. In the *second* stream, the *caregiver*’s face was blurred, and condition-specific audio cues were cut (**Supplementary Figure 1D**). For example, in the MO condition, the audio cues that follow “Go” in “Go, continue looking at your baby, and open your mouth” were cut out. Therefore, the experimental condition was masked (“blinded”). These “masked” videos were used (by separate coders) to code infants’ gestures.

### 1.4. Video coding and data quality checks

Test trial videos (n = 118, from 26 participants), with the *infant* face blurred (**Supplementary Figure 1C**), were evaluated based on whether video quality was sufficient for coding, and whether the participating caregiver engaged in the task and reached basic task fidelity criteria (i.e., modeling at least 4 target gestures in each 20s modeling period, and at most 0.2 target gestures in each 20s baseline/rest period, on average over the test trial). After these initial quality checks, 3 test trials (from 2 distinct dyads) were excluded due to insufficient video quality (e.g., lagging, freezing, poor resolution), 3 (from 3 distinct dyads) because the caregiver did not start the task (e.g., calming or feeding the child without modeling the gesture), and 16 (from 5 distinct dyads) were excluded because the caregiver did not reach task fidelity criteria (e.g., held their tongue out during the modeling period instead of repeatedly sticking it out and then retracting it, **Figure 1A**).

**Figure 1.**
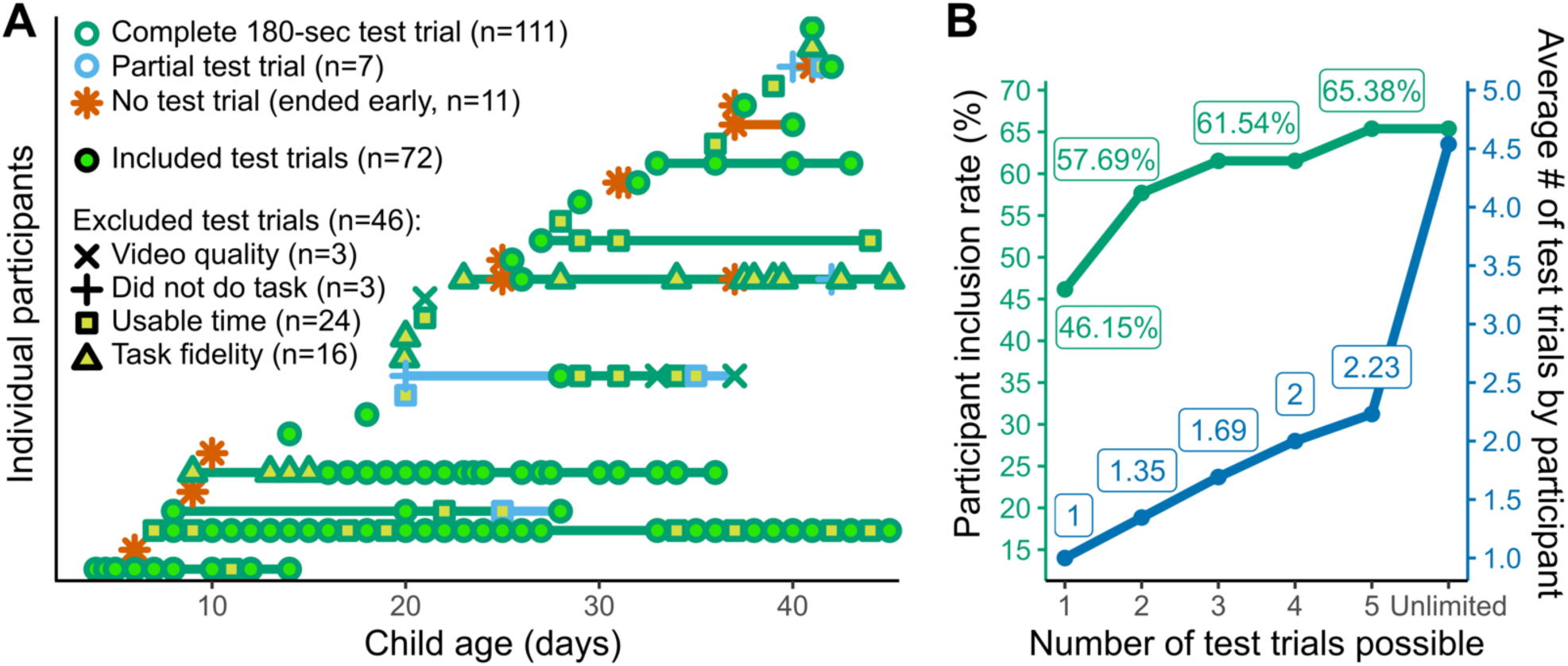
Repeated participation and data inclusion. **A.** N = 29 infants (4-45 days-old) and their caregiver participated in a total of n = 129 sessions, n = 118 of which yielded a complete or partial test trial, and n = 72 of which met inclusion criteria. Repeat participation was allowed, and 45% participated more than once. **B.** Allowing participants to contribute more than 1 test trial modestly raised the inclusion rate, with the largest increase observed when going from 1 to 2 test trials. Inclusion rates and average numbers of test trials are out of all participants who contributed at least one partial test trial (N = 26).

Test trials (n = 96, from 22 participants) that passed quality criteria were coded for infant gestures by separate team coders, this time with the *caregiver* face blurred and the infant face visible (**Supplementary Figure 1D**). TP was defined as a clear forward thrust of the tongue that recedes within 3 seconds, and MO was defined as a parting of the lips that reverses to a more closed position within 3 seconds (Heimann & Tjus, 2019). Frames were excluded as unusable if, for longer than 3 seconds, infants had their eyes closed, were crying, hiccupping, etc. A subset of test trials was coded by 2 independent coders to assess reliability (21 test trials; usable time: *r* = 0.91; mouth opening counts by 20s period: *r* = 0.83; tongue protrusion counts by 20s period: *r* = 0.83; Pearson’s *r*).

Finally, test trials were excluded if the test period had fewer than 80 seconds (out of 160) and/or the baseline period had fewer than 10 seconds (out of 20) of usable time (24 test trials, from 11 participants), resulting in a total of 72 included test trials from N = 17 participants (**Figure 1A**). Prior to analysis, infants’ MO and TP counts were converted to gesture rates per minute by dividing them by the corresponding amount of usable time, separately for both the initial 20s rest (“baseline”) period and for the 160s experimental period, and then multiplying by 60 (see **Supplementary Table 1**).

### 1.5. Session-level mimicry heuristic

Individual infants’ behavior varies across repeated sessions, motivating the definition of a session-level index of mimicry. To that end, we defined a session as showing evidence of mimicry (“imitator” session) if and only if both of the following two criteria are met: (1) there is a strictly positive increase in the rate of modeled gesture from baseline to test, <u>and</u> (2) this increase in rate from baseline to test is strictly higher for the modeled than for the non-modeled gesture (adapted from Paukner et al., 2017; Simpson et al., 2016). For example, in a session when a caregiver models MO, if an infant opens their mouth at a higher rate during test than baseline, and if that increase is larger than the increase in their rate of TP, then that session would be considered to show mimicry. For simplicity, we further refer to this heuristic as the “imitator” criteria.

### 1.6. Analysis

Analyses were conducted using custom code in MATLAB and R/RStudio. Bayesian statistics and associated simulations were run using R packages BayesFactor and BFDA (Rouder et al., 2017; Schönbrodt & Wagenmakers, 2018). Key analysis code is available at [masked].

## 2. Results

### 2.1. Paradigm feasibility and data quality

To assess the feasibility of conducting caregiver-led, asynchronous behavioral research with infants online in the first 6 weeks of life, we quantified the extent to which participating dyads completed the paradigm and contributed usable data.

#### Contributed test trials

Of 129 sessions from 29 participants (**Figure 1A**), 11 ended before the start of the test trial (“ended early”), i.e., without a test trial video. The remaining 118 sessions (from 26 participants) yielded test trial videos that were either complete (180s, n = 111 sessions) or incomplete (session ended after the start but before the end of the 180s test trial, n = 7 sessions). Thus, most sessions (91.5%), and most participants (90%) contributed at least a partial test trial. Of the 29 participating dyads, 13 (45%) initiated more than 1 session, and 9 contributed more than one test trial (31%; regardless of completion or inclusion). Thus, most participants were able to complete the paradigm at least partially, and almost 1/3 chose to participate in more than one session.

#### Inclusion and attrition

The resulting 118 test trials (61 MO and 57 TP, from N = 26 dyads) were coded for video quality and caregiver task fidelity, with participating infants’ faces blurred. Most test trials (81.4%) passed these data quality criteria. These 96 test trials (from N = 22 dyads) were separately coded for infant gestures. Of those, 24 test trials (from N = 11 dyads) were excluded due to having less than 10 seconds of usable time during the 20s baseline period, and/or less than 80 seconds of usable time during the 160s experimental period. The remaining n = 72 test trials (36 MO and 36 TP trials), from N = 17 participants, were included. Thus, the overall inclusion rate was 55.81% (72/129) at the session level and 58.62% (17/29) at the participant level. Focusing on sessions and participants with at least a partial test trial, the inclusion rate was 61.02% (72/118) at the session level and 65.38% (17/26) at the participant level. Thus, attrition was comparable to that reported in online asynchronous studies with older infants (Scott & Schulz, 2017) or in-person neonatal mimicry studies (Davis et al., 2021).

#### Impact of repeat participation on the overall participant inclusion rate

We next quantified the extent to which allowing repeat participation over subsequent days increased the proportion of participants who were able to contribute any included data, i.e., raised the overall participant-level inclusion rate. As expected, the participant-level inclusion rate increased modestly from 46.15% to 65.39% when allowing up to 5 repeat test trials compared to only the first — although with diminishing returns (**Figure 1B** ; inclusion rate increases from 46.15% to 65.39% when computed over N = 26 participants with 1 or more test trial, or from 41.38% to 58.62% when computed over all N = 29, in which case the denominator includes participants who initiated a session but did not attempt any test trial). No further gain in the participant inclusion rate was observed when allowing more than 5 repeat test trials, although repeat participation beyond that point may be of interest refine individual estimates or assess longitudinal change.

### 2.2. Participant demographics

Of the 29 participating caregivers, 26 identified as women (89.66%), and 3 as men. Participating caregivers were an average of 33.34 years-old (range: 23-55 years) overall and 34.59 years-old (range: 23-55 years) in the included sample, respectively. Because online asynchronous data collection has the potential to reach more families than laboratory-based studies, we assessed the extent to which our paradigm engaged a demographically representative set of US families. These data are summarized in **Figure 2**. Briefly, caregiver and family characteristics were broadly in line with comparison data (e.g., US Census) across all dimensions examined (PHQ-9 score, family ethnic/racial identity, household income, or geographic setting), except for significantly higher caregiver education levels compared to US Census data (**Supplementary Materials**). Included and excluded dyads did not statistically significantly differ by any of the demographic characteristics measured (infant sex, infant age at first session, caregiver age or gender, PHQ-9 score, education, family racial identity, household income, or geographic setting; all uncorrected *p*s > 0.05).

**Figure 2.**
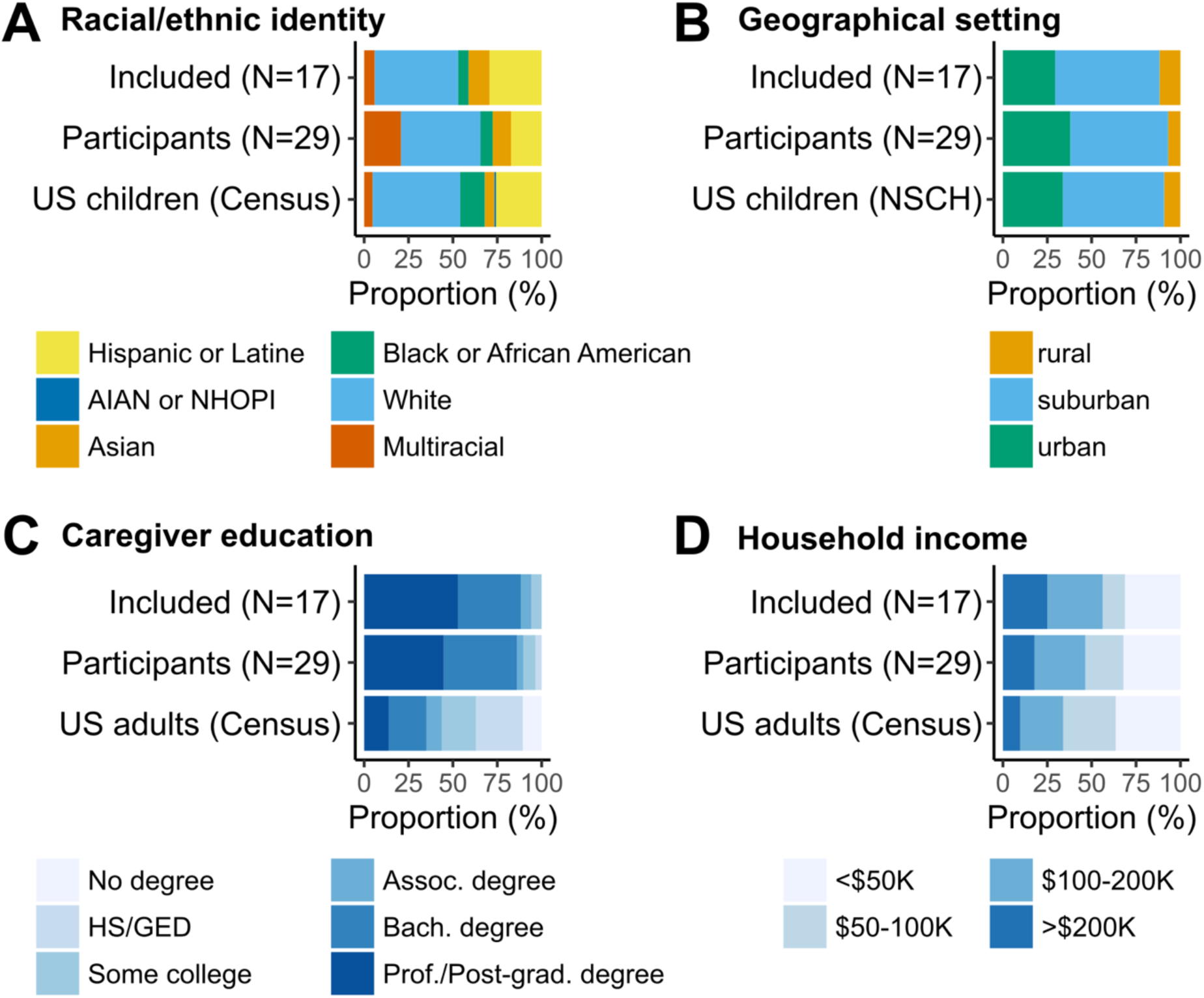
Demographic characteristics of participating infants and caregivers. **A.** Family race and/or ethnic identities. Hispanic or Latine includes all families identifying as such regardless of race(s). Other identities include only non-Hispanic or Latine families. AIAN: American Indian or Alaska Native; NHOPI: Native Hawaiian or Other Pacific Islander. **B.** Geographical setting. NSCH: National Survey of Children’s Health, from Calthorpe & Pantell (2021). **C.** Educational attainment of the participating caregiver. **D.** Household income.

### 2.3. Group-level mimicry of caregiver facial movements

We next sought to estimate the extent to which infants mimicked caregiver’s facial gestures in the current feasibility sample, to inform the design of better-powered future studies. Because different participants contributed different numbers of included sessions (i.e., 12 of 17 included participants contributed 1 included session, and the remaining 5 respectively contributed 3, 4, 9, 18, and 26 of a total of 72 included sessions), mimicry was analyzed both at the participant and at the individual session level.

#### 2.3.1. Participant-level analyses of neonatal mimicry

Of N = 17 included dyads, 8 caregivers were randomly assigned to model mouth-opening (MO), and 9 were assigned to model tongue-protrusion (TP). We used Bayesian and frequentist statistics to test whether infants produced more TP if their caregivers modelled TP rather than MO, and vice-versa. To derive participant-level estimates, rates of TP and MO were averaged over included sessions for each participant with more than one included session.

Infants’ rates of MO during the experimental period were not statistically significantly greater when caregivers modeled MO than when they modeled TP, although there was a marginal trend (M_MO condition_ = 9.52 gestures/min, M_TP condition_ = 6.54 gestures/min; d = 0.79; directional Bayes Factor: BF = 1.78; one-tailed *t*-test for independent samples, *t*[11.81] = 1.58, *p* = 0.070; **Figure 3A**). Infants’ rates of TP were not statistically significantly greater when caregivers modeled TP than when caregivers modeled MO, and there was moderate evidence for the null hypothesis (M_TP condition_ = 4.14 gestures/min, M_MO condition_ = 4.71 gestures/min; d = -0.23; BF = 0.32 < 1/3; *t*[14.49] = -0.47, *p* = 0.676; **Figure 3A**). Numerically comparable results were found when subtracting the baseline rate of each gesture (MOs: M_MO condition_ = 0.46 gestures/min, M_TP condition_ = -2.11 gestures/min; d = 0.50; BF = 0.96; t[14.71] = 1.05, *p* = 0.155; TPs: M_TP condition_ = -1.24 gestures/min, M_MO condition_ = -0.35 gestures/min; d = -0.16; BF = 0.34; *t*[13.30] = -0.33, *p* = 0.627; **Figure 3B**).

**Figure 3.**
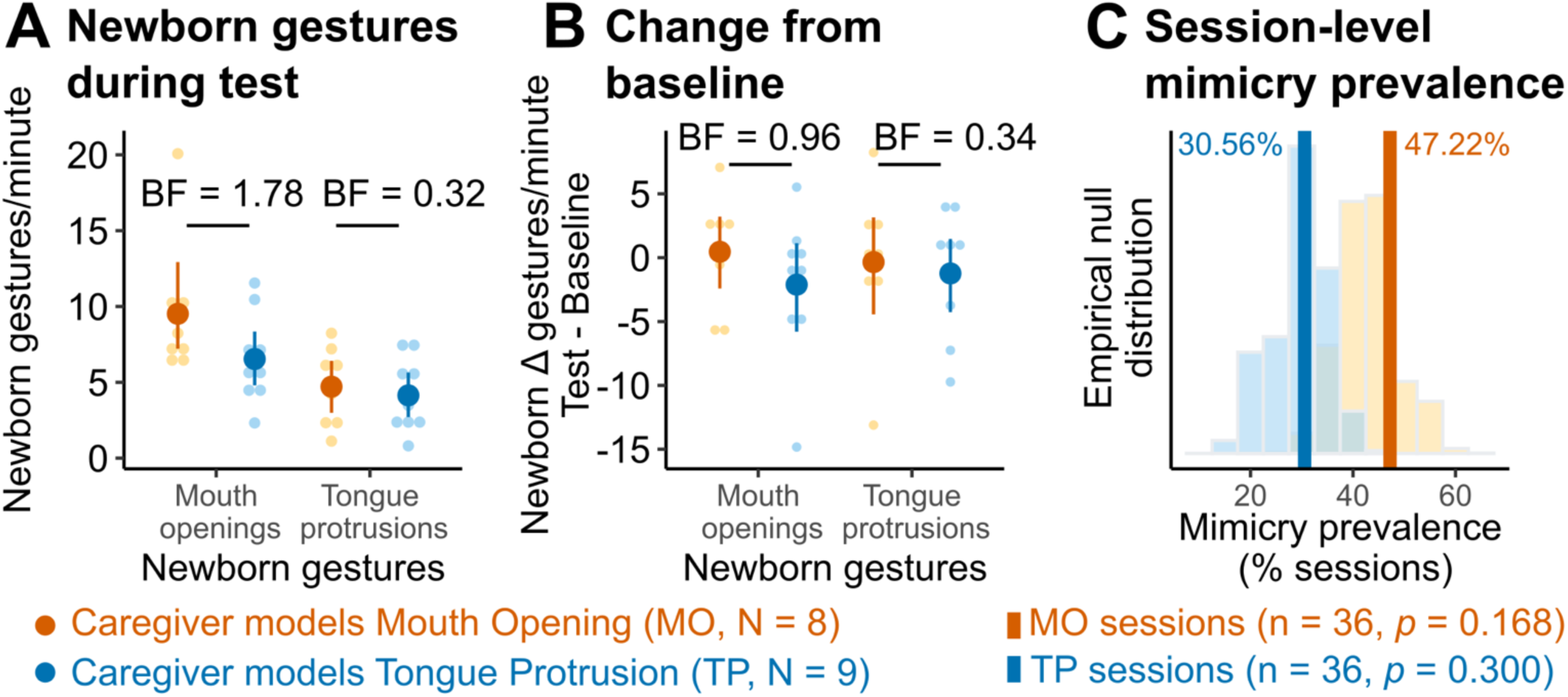
Limited to negative evidence of mimicry in the feasibility sample. **A.** Infants’ average rates of mouth openings or tongue protrusions were not significantly higher in response to matching caregiver movements. For tongue protrusions, there was moderate evidence favoring the null (directional Bayes Factor < 1/3). **B.** Correcting for baseline gesture rates yielded similar results. **C.** Session-level mimicry was not more frequent than expected by chance over the N = 36 sessions in either condition (“imitator” criteria, empirical p-values from 10,000 condition label permutations).

Overall, there was moderate evidence for the null hypothesis that infants’ average rates of TP were equal (or lower) in infants whose caregivers modeled TP than in infants whose caregivers modeled MO. In contrast, there was insufficient evidence to adjudicate whether infants’ average rates of MO were statistically higher in infants whose caregivers modeled MO than in infants whose caregivers’ modeled TP.

#### 2.3.2. Session-level analyses of neonatal mimicry

We next asked whether, when disaggregating infant behavior at the session level, more sessions showed behavior consistent with mimicry according to the “imitator” criteria than expected by chance (adapted from Paukner et al., 2017; Simpson et al., 2016; see **Methods**). To that end, we derived empirical null distributions, i.e., computed the proportion of MO and TP modeling sessions that would be considered to show mimicry if all condition labels were randomly shuffled (10,000 permutations), then compared the observed proportions of “imitator” sessions observed in the data to these null distributions.

Caregivers modeled mouth opening (MO) in 36 of the 72 included sessions, and tongue protrusion (TP) in the remaining 36 included sessions. As defined by the “imitator” criteria, infants showed evidence of possible mimicry in 47.22% (17/36) of included MO sessions and 30.56% (11/36) of TP sessions. These proportions of MO and TP sessions showing possible evidence of mimicry did not statistically significantly differ from what would be expected by chance, i.e., if condition labels were randomly shuffled (empirical one-sided p-values; MO: *p* = 0.168; TP: *p* = 0.300; **Figure 3C**).

#### 2.3.3. Projected sample size to reach conclusive evidence

To inform potential future studies, we used simulations to estimate the sample size needed to reach a more conclusive level of evidence. Specifically, we asked how many participants would need to be included to reach a target Bayes Factor (BF) of less than 1/6 or higher than 6, in a sequential Bayesian, between-participants design (Schönbrodt & Wagenmakers, 2018; Stefan et al., 2019), separately comparing infants’ average gestures rates in each condition with directional Bayesian tests and no correction for multiple comparisons across the two gestures.

If the true effect size for a given gesture was as large as that observed for MO in the current study (d = 0.79), then with a maximum total included N = 300 (150 by group), 100% of studies would conclusively reject the null (BF of 6 or larger), with a 0% false negative rate. On average, studies would terminate at a total sample size of N = 68 (34 by group), with 80% terminating at N = 66 or lower. Under the null hypothesis (that is, if the true effect size is equal to or lower than 0, as observed for TP in the current study), 80.5% of studies would reach a conclusive level of evidence for the null (BF of 1/6 or smaller), with a 4.3% false positive rate. Such studies would terminate at an average total N = 138, with 80% of studies terminating at N = 240 or lower. Thus, in either case, a maximum included N = 300 would be sufficient to reach conclusive evidence for the ground truth in at least 80% of studies, and at least 80% of studies would terminate with at N = 240 or fewer included participants.

We next estimate how many test trials would need to be collected to reach N = 300 maximum included participants, given the attrition and participation patterns observed in the current study (**Figure 1B**). Allowing up to 2 repeat test trial completions by participant was sufficient to increase the inclusion rate past 50%. Based on that observed inclusion rate for up to 2 repeat test trial completions allowed, at least one test trial (partial or complete, included or not) would need to be collected from N ∼ 520 participants to reach N = 300 maximum participants included. We would expect that each would contribute an average of ∼1.35 test trials, leading to a maximum projected total of n ∼ 700 test trials contributed. By comparison, the current study collected a total of n = 118 test trials. Therefore, the current paradigm would need to be scaled up at most ∼6 times (test trials-wise) to reach a conclusive level of evidence (BF < 1/6 or BF > 6) in at least 80% of cases under either ground truth scenario outlined above. A within-participant design would reach adequate statistical power at a lower sample size, though potentially at the expense of higher attrition or cross-conditions spillover.

### 3.4 Caregivers’ judgments of neonatal mimicry

For neonatal mimicry to plausibly promote dyadic bonding or interactions, caregivers need to be able, implicitly, or explicitly, to perceive neonatal mimicry when it occurs—regardless of whether such perceived mimicry is genuine, artefactual, or coincidental. As an initial foray into this question, we asked whether caregivers’ binary perception of mimicry in each session aligned with the session-level index of mimicry as described above (“imitator” criteria). Surprisingly, caregivers’ binary reports of mimicry in individual sessions did not significantly align with this session-level, researcher-defined index of mimicry, i.e., detectability indices did not significantly differ from their theoretical chance level of 0 (d’, **Table 1**). Compared to the researcher-defined index, caregivers in the current sample also tended to be liberal in reporting TP mimicry (i.e., c-bias < 0; 83% of all and 75% of first TP sessions reported as imitative), but conservative in reporting MO mimicry (i.e., c-bias > 0; 25% of all and 22% of first MO sessions reported as imitative; **Table 1**).

**Table 1.** Signal detection metrics comparing caregivers’ reports with a standard definition of mimicry at the session level (see main text). Point estimates and 95% CIs derived from 5000 bootstrapped samples. MO: Sessions in which caregivers modeled mouth opening. TP: Sessions in which caregivers modeled tongue protrusion.

|  | Sessions | Detection<br>index (d') | Conservative<br>bias (c-bias) | Area Under the<br>Curve (AUC) |
| --- | --- | --- | --- | --- |
| All included<br>sessions | 72 | -0.16<br>[-0.75, 0.46] | -0.08<br>[-0.39, 0.22] | 0.46<br>[0.32, 0.60] |
| MO only | 36 | -0.08<br>[-1.00, 0.80] | 0.64*<br>[0.19, 1.07] | 0.46<br>[0.29, 0.66] |
| TP only | 36 | 0.35<br>[-0.90, 1.13] | -0.98*<br>[-1.38, -0.37] | 0.58<br>[0.36, 0.77] |
| First<br>included<br>session | 17 | 0.11<br>[-1.20, 1.37] | 0.05<br>[-0.61, 0.68] | 0.58<br>[0.21, 0.88] |
| MO only | 9 | 0.15<br>[-1.38, 1.53] | 0.60<br>[-0.41, 1.25] | 0.55<br>[0.00, 0.90] |
| TP only | 8 | 0.60<br>[-0.59, 1.80] | -0.67*<br>[-1.28, -0.09] | 0.83<br>[0.31, 1.00] |
| Longitudinal<br>participants | 53 | -0.21<br>[-0.91, 0.49] | -0.04<br>[-0.41, 0.31] | 0.42<br>[0.25, 0.59] |
| Theoretical chance level |  | 0 | 0 | 0.5 |

Caregivers additionally reported how confident they felt about each binary judgment. In 90% of included sessions (and in 100% of the first included attempts), caregivers reported at least a mild level of confidence in their binary report (i.e., 3 or more on a scale of 0-5). Caregivers’ binary reports of mimicry were combined with their confidence reports to derive scores ranging from -5 (high confidence of no mimicry) to +5 (high confidence of mimicry; **Figure 4**). Pooling these scores across all sessions allowed for the additional derivation of group-level ROC curves, further quantifying the alignment of caregivers’ perceptions with the researcher-defined index of neonatal mimicry in each session. The resulting Area Under the Curve (AUC) provided a group-level metric of the alignment between caregivers’ perceived mimicry in each session and whether a standard researcher-defined index (as described above) would classify infants’ behavior in that session as imitative. Again, the resulting AUCs did not significantly differ from chance (AUC, **Table 1**), i.e., these two measures did not align. Similar results were obtained when restricting analysis to the first included session of each dyad, or to sessions from only those 3 “longitudinal” participants with 9 or more included attempts each (**Table 1** and **Figure 4AB**).

**Figure 4.**
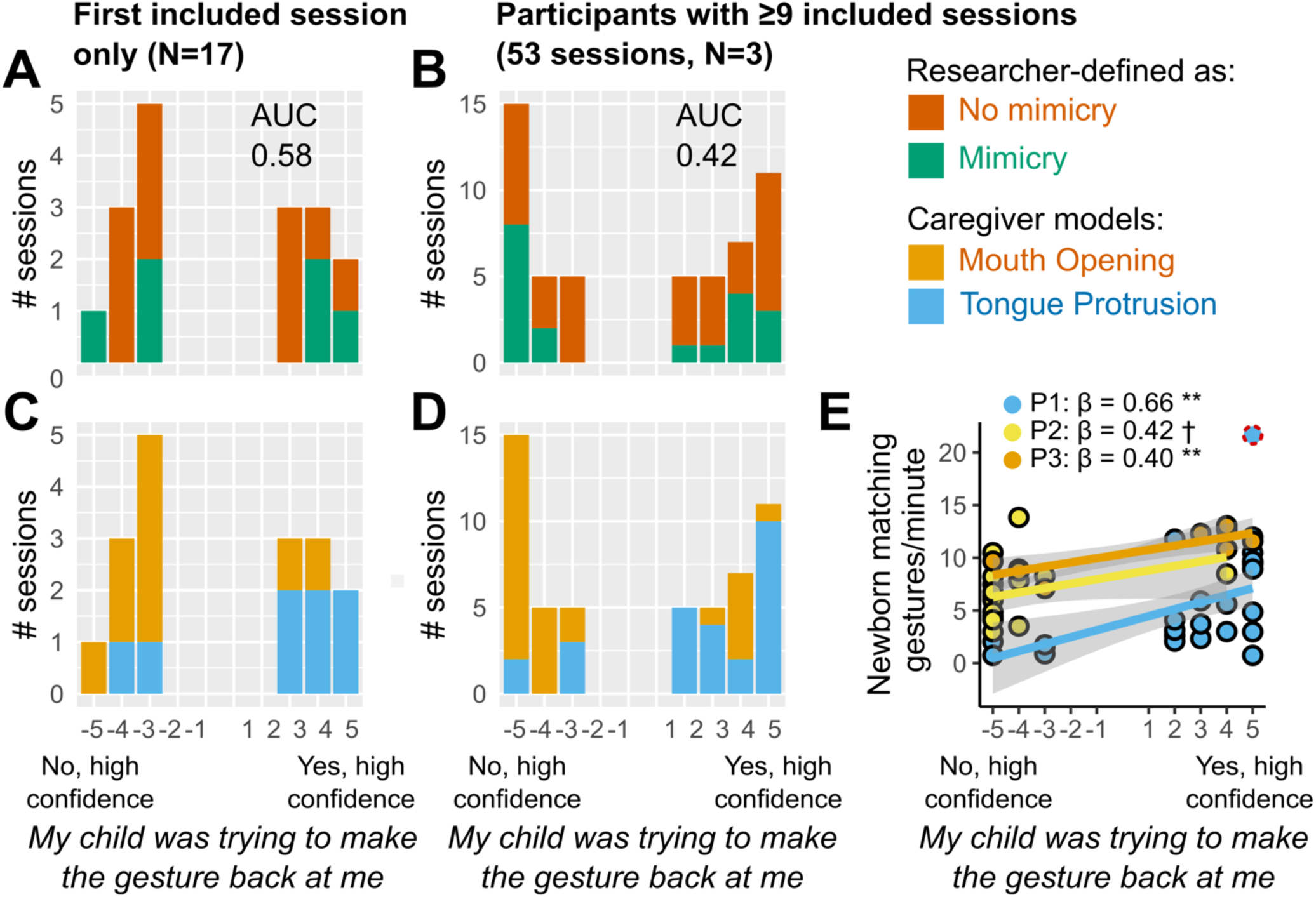
Caregiver perceptions of mimicry in individual sessions. After each session, caregivers reported whether they thought their child was trying to mimic them (forced choice), and their confidence in their answer (5-point Likert scale). Considering either the first included session only for all participants (**A**, **C**), or only longitudinal participants with 9 or more included sessions (**B**, **D**): **A**, **B**. Caregivers’ reports did not align with a standard session-level definition of mimicry. **C**, **D**. Caregivers reported more mimicry if they modeled tongue protrusion. **E**. “Longitudinal” caregivers tended to report more mimicry in those sessions when their child made more matching gestures (N = 3 participating dyads noted as P1, P2, and P3). Individual linear fit lines: † *p* < 0.1, * *p* < 0.05, ** *p* < 0.01 (FDR-corrected over the 3 fits). Outlier datapoint (red dotted outline) excluded from corresponding linear fit.

Taken together, these results suggest that caregivers’ reports of whether their child was trying to mimic them may reflect different factors than what the session-level heuristic above captures. A possibility is that caregivers are sensitive to their infants’ matching gestures, even if they aren’t as sensitive to “control” variables (baseline rate of matching gesture, non-matching gesture rate) as the standard index is. To explore this possibility, we next tested whether higher infant matching gesture rates (e.g., rates of mouth opening if the caregiver modeled mouth opening) were associated with higher caregiver detection scores, controlling for the experimental condition (mouth opening vs. tongue protrusion modeling). Across all included sessions, caregivers reported more mimicry (i.e., detection report scores were higher) when they had been randomly assigned to model tongue protrusion compared to mouth opening (linear mixed-effect model with random intercept and slope, χ^2^[1] = 52.49, *p* < 0.001), consistent with the conservative bias findings above. Caregivers also reported more mimicry when their infants made more matching gestures (χ^2^[1] = 4.80, *p* = 0.029), suggesting a sensitivity of caregivers’ reports to infants’ behaviors. The effect of the experimental condition remained significant when considering only the first included session for each dyad (linear regression, β = 5.30, *F*[1,13] = 9.81, *p* = 0.008, 1 outlier removed with Cook’s distance > 1; **Figure 4C**). The effect of infants’ matching gestures rate remained numerically evident, although only marginally significant, in that subset (linear regression, β = 0.62, *F*[1,13] = 4.36, *p* = 0.057). The latter effect was also evident in the 3 “longitudinal” dyads who contributed 9 or more included sessions each (total of 53). This association between caregivers’ reports and infants’ matching gesture rates also remained present, either marginally or significantly, when individually considering each of the “longitudinal” dyads (N = 3 dyads, total of 53 sessions, **Figure 4E**).

## 3. Discussion

The present study demonstrates the feasibility of engaging US families of young infants (0-6 weeks old) in asynchronous online behavioral research. There was moderate evidence that infants in this sample did not mimic their caregivers’ tongue protrusions. In addition, there was ambiguous evidence on whether they mimicked their caregivers’ mouth openings. Caregivers’ perception of mimicry in individual sessions was associated with infants’ matching gesture rates, but not with a standardized definition of mimicry.

To our knowledge, the current study is the first to successfully engage caregivers and 0-6 weeks-old infants in online, asynchronous, webcam-based research. Most dyads were able to complete the paradigm at least partially, and almost a third chose to participate more than once. Allowing just one repeat session (i.e., up to 2 sessions by dyad) sufficed to reach an inclusion rate comparable to that of online studies with older infants (Scott & Schulz, 2017), or of in-person neonatal mimicry studies (Davis et al., 2021). Streamlining and clarifying task instructions for caregivers participating asynchronously (e.g., more explicit guidance on target speed when modeling face movements, shorter practice) may improve attrition further. Substantial variability in the timing and number of repeat sessions constrained statistical analyses of the resulting data, though it maximized caregivers’ ability to participate when available. Future studies may thus encourage a specific target number and timing of sessions: for example, aiming for 2-3 sessions in a short timeframe to boost participant inclusion in single age-group, or for repeated longitudinal sessions to investigate learning or developmental change. While online webcam-based studies can present challenges for families who lack access to broadband Internet (Sheskin et al., 2020), the current sample roughly matched US demographics in terms of geographic area, race, and ethnicity. To better represent US families, additional efforts are needed to engage caregivers without a 4-year college degree. Data collection in online asynchronous research being inherently more scalable than in-person research, future studies may utilize this medium to collect data from larger samples of infant-caregiver dyads. Nonetheless, processing and hand-coding of infant behavior videos remain a significant bottleneck to scalability, due to the current absence of widely available, validated, and fully automated pipelines for coding relatively subtle infant movements such as tongue protrusion (although, for MO, see Zeng et al., under review). Despite this limitation, the current findings support the feasibility of leveraging asynchronous online testing to broaden the participation of newborns and young infants in psychological research, where they remain under-represented.

To inform potential future studies, we assessed whether infants showed evidence of mimicking their caregivers’ gestures in this feasibility sample of 72 sessions from N = 17 dyads. Specifically, we asked whether infants’ average rates of MO were higher if their caregivers modeled MO than if their caregivers modeled TP, or vice-versa. There was no clear evidence for the mimicry of mouth opening, and moderate evidence against the mimicry of tongue protrusion. Similar results were found when estimating mimicry rates with a session-level heuristic (adapted from Simpson et al., 2016). Simulations based on effect sizes estimated in the current sample further suggested that scaling the current design by a factor of up to ∼6, that is, collecting ∼700 test trials to include up to N = 300 participants (sequential Bayesian, between-participants design) is expected to reach a conclusive level of evidence (Bayes Factor >6 or < 1/6). Materials necessary to replicate or build upon the current paradigm are openly shared online to facilitate future research.

It is possible that, in addition to the relatively small sample size in a single-trial between-subjects design, the use of an asynchronous online medium (Chuey et al., 2024; Raz et al., 2024), with caregivers as models, and in the inherently “noisy” home environment, contributed to the present negative findings. However, modeling by caregivers in the home, without synchronous supervision by an experimenter, arguably makes the current findings more applicable than lab-based findings with experimenters as models to the question of whether newborns and young infants mimic face movements in spontaneous, real-life social interactions, as these interactions disproportionately occur with caregivers and under more variable conditions than the laboratory environment. In addition, the current findings are in line with a previous in-person study (N = 11, 14-21 days-old) of young infants at home, which also focused on caregivers’ mouth opening and tongue protrusion, and did not report statistically significant mimicry (Heimann & Schaller, 1985). One other study with mothers as models in a hospital setting also found that newborns (N = 18, 35-68 hours-old) were not statistically more likely to produce mouth openings in response to mouth openings than in response to tongue protrusions, or vice versa (Ullstadius, 1998). Another study using mothers as models in a strictly controlled laboratory procedure (N = 32, 6-weeks-old) did find evidence of mimicry for tongue protrusions, but not for mouth openings (Meltzoff & Moore, 1992). In other words, while the current sample size was small, the findings generally align with previous mixed or negative evidence of neonatal mimicry when using caregivers as models. If confirmed by larger studies, such negative finding would argue against a general (i.e., group-level) role of mimicry in social visual learning and bonding during the first few postnatal months, as these processes predominantly occur in interactions with caregivers (Jayaraman et al., 2015; Sugden et al., 2014). Cross-modal representations of facial movements may thus generally emerge closer to 5 months of age, as both sensorimotor representations of one’s face, visual acuity, and high-level visual representations of others’ faces sharpen (Isomura & Nakano, 2016; Kobayashi et al., 2014).

For neonatal mimicry to potentially boost caregivers’ bonding or propensity to engage in face-to-face expression play (Duffy & Chartrand, 2015; Meltzoff & Marshall, 2018) caregivers need to be able to (at least implicitly) detect the occurrence of mimicry during face-to-face interactions with their infant. For example, if only some individual young infants tend to overtly mimic gestures (Heimann & Schaller, 1985; Kaburu et al., 2016; Simpson et al., 2016), then do their caregivers, implicitly or explicitly, detect those instances? Or, if neonatal mimicry is artefactual (Keven & Akins, 2017), then do caregivers nonetheless tend to interpret those instances as evidence of mimicry? As a starting point to addressing these questions, we asked caregivers to report whether they thought their child’s behavior during each test trial was indicative of mimicry. Caregivers’ reports of mimicry did not align with an often-used session-level heuristic of mimicry (imitator vs. non-imitator, Paukner et al., 2017; Simpson et al., 2016). This mismatch suggests that more research is needed to capture aspects of infants’ behavioral responses that caregivers generally perceive as indicative of mimicry (e.g., see also Heimann & Schaller, 1985). As expected, caregivers’ perceptions of mimicry were positively associated with infants’ matching gesture rates. Surprisingly, caregivers were also more likely to ascribe mimicry to their infant if they were randomly assigned to model tongue protrusion rather than mouth opening. Caregivers might have perceived tongue protrusions as less likely to occur by chance and, therefore, have been more inclined to interpret them as imitative. Alternatively, caregivers may have been influenced by prior exposure to common depictions of neonatal mimicry showing tongue protrusion (such as in popular media or textbooks).

In conclusion, the current study provides an initial proof-of-concept for the feasibility of using an open-source platform to conduct asynchronous online behavioral research with young infants and their caregivers. Initial findings leaned towards the null hypothesis that these infants generally (i.e., at least at the group level) do not mimic their caregivers’ tongue protrusions and failed to find support either for or against infants’ mimicry of their caregivers’ mouth openings. Simulations suggest that larger sample sizes (e.g., maximum target N = 300) are needed to reach a more conclusive degree of evidence. Caregivers’ perceptions of mimicry reflected infants’ matching behaviors but did not align with an often-used mimicry index, suggesting that more research is needed to capture behaviors that caregivers interpret as imitative. Materials and data are shared to facilitate replications or extensions of the design. Future research may build upon and scale up the current online design to achieve stronger levels of evidence at the group level, examine longitudinal effects or developmental changes from 0 to 6 weeks, obtain robust individual estimates, or further investigate factors that shape caregivers’ perceptions of mimicry in interactions with their infants.

### Data Availability

Processed data and materials are available at [masked]. Raw videos are shared to authorized users on Databrary, according to caregiver privacy choices, at http://databrary.org/volume/1300/.

## Acknowledgements

This work was partly supported by an NSF CAREER Award 1653737 and an APS James McKeen Cattell Fund Fellowship Sabbatical Award to EAS. LB acknowledges the support of a Young Investigator Award from the Brain & Behavior Research Foundation, a Karen Toffler Charitable Foundation award, and a CAS Collaborative Pilot grant from American University. We thank the participating families, research assistants, and students who made this work possible, and Kimberly M. Scott for feedback and essential technical support in designing and implementing the study on Children Helping Science (formerly known as Lookit). The authors declare no competing financials interests.

## Author Contributions

Conceptualization & Supervision, E.A.S and L.B.; Methodology, E.A.S and L.B.; Software, K.C., Z.G., K.A., E.A.S, L.B.; Formal Analysis & Visualization, K.C., J.G., L.B.; Investigation & Data Curation, K.C., C.L., Z.G., E.A.S, and L.B.; Writing – Original Draft, B. N., K.C., L.B.; Writing –Review & Editing, B. N., K.C., C.L., Z.G., K.A., J.G., E.A.S, and L.B..

